# Is the winter survival area of *Empoasca fabae* continuously expanding?

**DOI:** 10.64898/2026.05.25.727670

**Authors:** Abraão Almeida Santos, Edel Pérez-López

## Abstract

*Empoasca fabae* is a migratory pest that overwinters in the Southeastern United States (US) and damages crops throughout its summer range in North America. Its spring arrival has advanced by 9.7 days between 1951 and 2012, and increased damage is linked to warmer conditions that accelerate host and pest development. Yet whether this advance is also associated with a northward expansion of its winter survival area remains an open question.
Here, we analyzed 126 years of minimum temperature data across the contiguous US to address this question. Using its winter survival threshold (−9°C), we calculated the annual winter survival area for *E. fabae* (temperature-only) and tested for time-series trends. We also mapped the potential overwintering area (temperature + winter hosts) under evergreen and pine-only forest scenarios.
The estimated winter survival area varies over 126 years, showing a nonlinear pattern. However, we found no significant trend, change-point year, or rate of change. This lack of significance was also observed when considering the 1951–2012 period.
The Southeastern US remained consistently suitable for winter survival, while the northern edge varied latitudinally, especially within the range of 35°–40°N, with no clear trend.
Potential overwintering areas extend into Central Florida and the Texas Gulf Coastal Plain but exclude parts of Tennessee. In the pine-only scenario, the area in Mississippi and Alabama would be smaller.
The winter survival area for *E. fabae* has not continually expanded. The Southeastern US area remains suitable for over 126 years, whereas the northern range varies dynamically.

**GRAPHICAL ABSTRACT AND HIGHLIGHTS:** 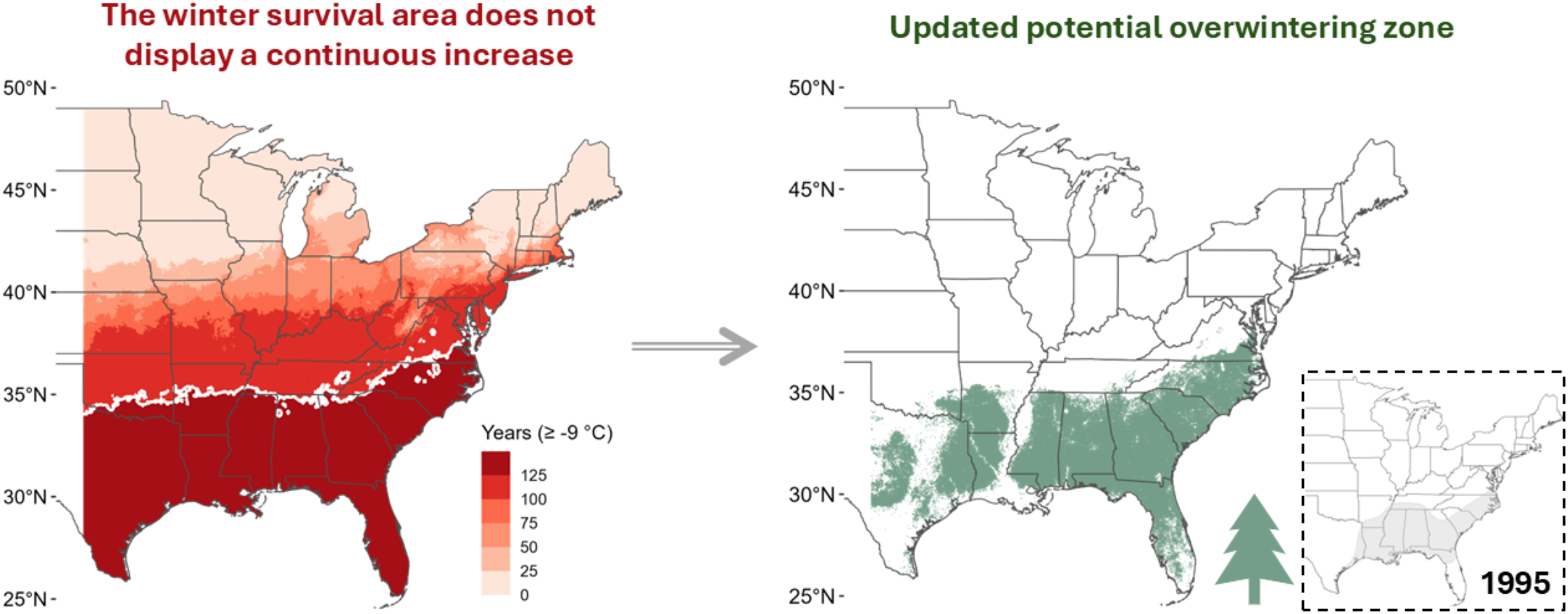

- The winter survival zone (temperature only) of *Empoasca fabae* has not expanded continuously over the past 126 years.
- The Southeastern United States remained suitable over this period, with the maximum northward extent of potential survival reaching 45°N and high variation within the range of 35°–40°N.
- Updated potential overwintering zones (temperature + winter hosts) extend into Central Florida and the Texas Gulf Coastal Plain.

## INTRODUCTION

*Empoasca fabae* (Harris) (Hemiptera: Cicadellidae) is a polyphagous leafhopper pest native to North America (DeLong, 1938; Lamp et al., 1994; Ross et al., 1964). Each spring, this species undertakes long-distance migrations from its estimated overwintering area in the Southeastern United States (US) (Carlson et al., 1992; Decker & Cunningham, 1968; Taylor & Shields, 1995) to the Midwest and the Northeastern US (Baker et al., 2015; Maredia et al., 1998), as well as Eastern Canada (Plante et al., 2024; Saguez et al., 2014). Its summer distribution appears to be limited to the west of 100° longitude (DeLong, 1938), although occasional reports may occur outside this limit (Chasen et al., 2014). By fall, they return to the overwintering area, where evergreen forests (especially pines) serve as the primary winter hosts (Decker & Cunningham, 1968; Taylor et al., 1993; Taylor & Shields, 1995; Shields & Testa, 1999).

The ongoing temperature increases have advanced the spring arrival of *E. fabae* by 9.7 days (from 1951 to 2012), but more significantly, worsening its impact on crops across the US (Baker et al., 2015). This phenomenon is thought to be associated with: *(i)* the acceleration of host development; *(ii)* a shortened *E. fabae* development time; and *(iii)* potential changes in the spring synoptic wind system, although this last point is less well understood (Baker et al., 2015). Nevertheless, could these early arrivals and subsequent increased damage also be linked to a continuously expanding winter survival zone?

The overwintering area for *E. fabae* was estimated spatially in 1995 (Taylor & Shields, 1995) by combining seasonal field surveys across the Southeastern US with leafhopper detections on evergreen plants, especially pines, resulting in a larger area in this southern region than previously thought (Decker & Cunningham, 1968). Then, in 2005, Sidumo et al., (2005) refined this estimate using a −9°C threshold, suggesting that the overwintering area could be extended farther north than in the 1995 estimate. Given these variations in the estimated overwintering area and the related advance and crop damage (Baker et al., 2015), we asked: Is the winter survival zone for *E. fabae* continually expanding? If so, is there a specific year that marks a turning point in this change?

Here, we analyzed 126 years (1900-2025) of spatial minimum temperature data and conducted a time-series analysis of the estimated winter survival area to answer these questions. We were also interested in identifying the current potential overwintering area, given the main winter hosts of *E. fabae*. From now on, by winter survival area, we mean the spatial extent where winter temperatures remain at or above −9°C (Sidumo et al., 2005). By overwintering area, we refer to the intersection of this winter survival area and the presence of *E. fabae* winter hosts (Decker & Cunningham, 1968; Taylor & Shields, 1995).

## METHODS

### Spatial data acquisition and raster analyses

To analyze the *E. fabae* winter survival area, we retrieved monthly minimum temperature data (1900–2025) for the contiguous US at a spatial resolution of 800 m from the PRISM database (PRISM Climate Group, 2025). For each year, we identified the colder month between January and February and extracted the corresponding raster (n = 126 raster files). By May 2026, all the rasters used in our study were classified by PRISM as ‘unlikely to change’ at the time of analysis.

We estimated the winter survival area of *E. fabae* at its summer longitudinal limit (100°W) (DeLong, 1938) using a minimum-temperature survival threshold of −9°C (Decker & Maddox, 1967; Decker & Cunningham, 1968), following Sidumo et al., (2005). We restricted our analysis to 100° W, consistent with DeLong (1938), who indicated this as the limit of *E. fabae’s* summer distribution. Although occasional collections have occurred farther west, e.g., in California (Chasen et al., 2014), the main distribution and economic impact zone lies on the 100°W boundary (Baker et al., 2015).

For each year, raster cell data were converted to binary using the temperature threshold, with 0 indicating areas below −9°C and 1 indicating areas at or above −9°C. We then estimated the winter survival area (km^2^) by summing the areas of raster cells classified as 1. Subsequently, we combined the 126 binary rasters to visualize spatial variation in the survival area over time. Using the common winter survival area across all 126 years (i.e., the area where all cells remained as 1), we generated a raster file with this information to further estimate the overwintering area.

We considered two scenarios to estimate the overwintering area: evergreen forests and pines only, given that these hosts are thought to be the primary ones on which *E. fabae* feeds during winter (Taylor et al., 1993; Taylor & Shields, 1995). Evergreen data were obtained from the USDA Cropland Data Layer 2024 at 30 m resolution (Crop Land Data, 2025), while pine data were obtained from the 2024 Existing Vegetation Type (EVT) dataset at 250 m resolution (LANDFIRE, 2025). We selected EVT categories using keywords related to pines [loblolly, longleaf, P(p)inus, P(p)ine, slash, shortleaf] and combined them into a single layer. Both datasets were cropped to match *E. fabae*’s longitudinal extent and resampled to 800 m to align with the temperature data. We initially combined the rasters to visualize the overwintering area, then overlaid them to filter out regions without forest within the survival area. Finally, we calculated the area (km^2^) of each forest type’s overwintering zone and converted the rasters to polygons for visualization.

### Statistical analyses

All analyses and visualizations were conducted in R (version 4.4.2) and RStudio (version 2024.12.1), using the ggplot2 package (Wickham, 2016), the terra package for raster analyses (Hijmans, 2025), and the US map from the rnaturalearth package (Massicotte & South, 2025).We were interested in whether there is an upward trend in the winter survival area for *E. fabae*. Then, we first analyzed the annual time series (1900-2025) of the estimated winter survival area (km^2^) and, for comparison, the 1951-2012 series (Baker et al., 2015). Prior to trend analysis, we assessed time-series autocorrelation using the Ljung-Box test (lag = 10), the Augmented Dickey-Fuller (ADF) test for unit roots, and the Kwiatkowski-Phillips-Schmidt-Shin (KPSS) test for stationarity. The time series were stationary [(1900-2025 ADF: p = 0.01, KPSS: p = 0.1); (1951-2012 ADF: p = 0.01, KPSS: p = 0.1)] and showed no significant autocorrelation [(1900-2025: χ^2^ = 5.57, df = 10, p = 0.85); (1951-2012: χ^2^ = 4.24, df = 10, p = 0.93)].

To visualize the underlying pattern while smoothing short-term variability, we applied locally estimated scatterplot smoothing (LOESS) with a smoothing span of 0.75. We tested for monotonic trend using the Mann-Kendall test (α = 0.05; Kendall package; McLeod, 2022) and quantified the rate of change using Sen’s slope estimator (trend package; Pohlert, 2023). Finally, we applied Pettitt’s test (α = 0.05; trend package; Pohlert, 2023) to identify potential change points in the series.

## RESULTS

The estimated winter survival area varied over 126 years, displaying a non-linear trend in the calculated area (**Figure 1a**). Initially, the area tended to increase, then declined by the 1950s, with a slight upward reappearing from 1980 to 2025 (**Figure 1a**). However, we found that this trend was not significant (Mann-Kendall test: τ = 0.02; two-sided p-value = 0.67), nor was the estimated rate of change [Sen’s slope: z = 0.42; n = 126; slope (95% CI) = 403.98 (−1539.47 to 2570.64); p-value = 0.67]. Although Pettitt’s test indicated a change point around 1986, it was not significant either (U* = 534; p-value = 0.85).

**FIGURE 1.**
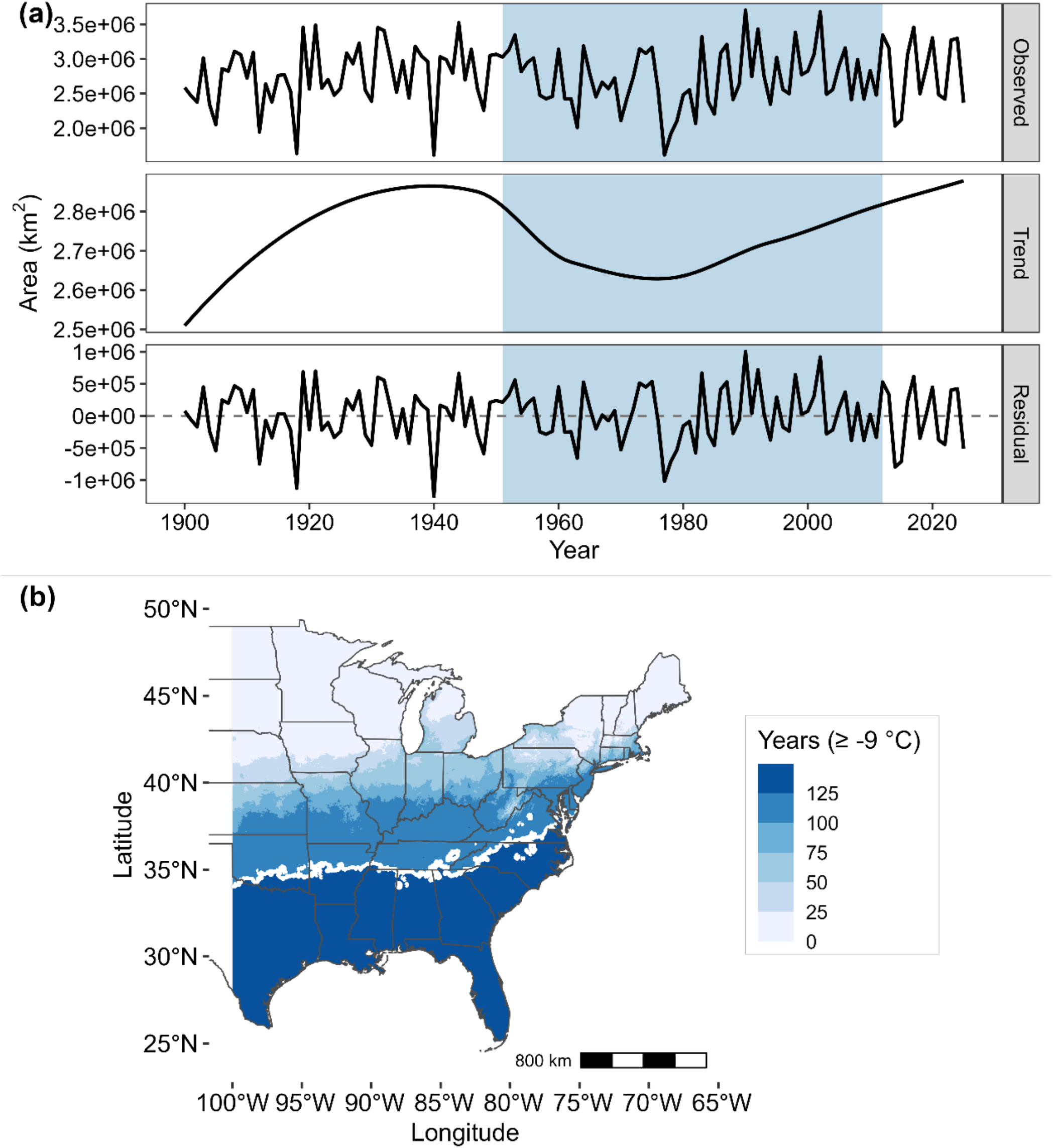
Estimated winter survival area (km^2^) for *Empoasca fabae* from 1900 to 2025. In (a), panels show observed values, the LOESS-smoothed trend component (span = 0.75), and residuals representing year-to-year variability. The winter survival area was calculated using the −9°C survival temperature threshold and restricted to longitudes west of 100°W. The blue-highlighted region (1951-2012) denotes the study period from Baker et al., (2015). In (b), the annual spatial frequency of the estimated winter survival area is shown. The white isoline marks the area where all years overlapped.

In our second time-series analysis, covering 1951 to 2012, which corresponds to the period studied by Baker et al., (2015), the results showed no evidence of a significant trend (τ = 0.06; two-sided p-value = 0.48) and no significant rate of change [z = 0.70; n = 62; slope (95% CI) = 1580.23 (−3512.70 to 8510.69); p-value = 0.48]. Similarly, Pettitt’s test indicated a change point around 1986, but it was not significant (U* = 287; p-value = 0.26).

The extension of the winter survival area remained suitable along the longitudinal boundary of *E. fabae* occurrence (100°W), with variations in the latitudinal range (**Figure 1b**). The maximum northern extent reached up to 45°N, but such variations were less common at this latitude. Conversely, there was a high frequency of years in which the winter survival area extended to 35°–40°N. Interestingly, over 126 years, the winter survival zone consistently included at least the Southeastern U.S., with an estimated area of 1,406,735 km^2^ (**Figure 1b**, white isoline).

We confirmed the potential overwintering area for *E. fabae* in the Southeastern US, as proposed in 1995 (Taylor & Shields, 1995) (**Figure 2a-e**). The overwintering area based on evergreen forests, with an estimated area of 299,047.5 km^2^ (**Figure 2b-c**), was larger than that for pine forests, with an estimated area of 65,905.9 km^2^ (**Figure 2d-e**). Both current estimates suggest a slight expansion of the overwintering region into Central Florida and the Gulf Coastal Plains of Texas (**Figure 2c-e**) compared to the initial estimate (**Figure 2a**). Meanwhile, areas in Tennessee seem less suitable due to the lack of winter hosts. If pine forests are the only winter hosts, this could result in a smaller overwintering area in Mississippi and Alabama (**Figure 2e**) compared to evergreen forests (**Figure 2c**).

**FIGURE 2.**
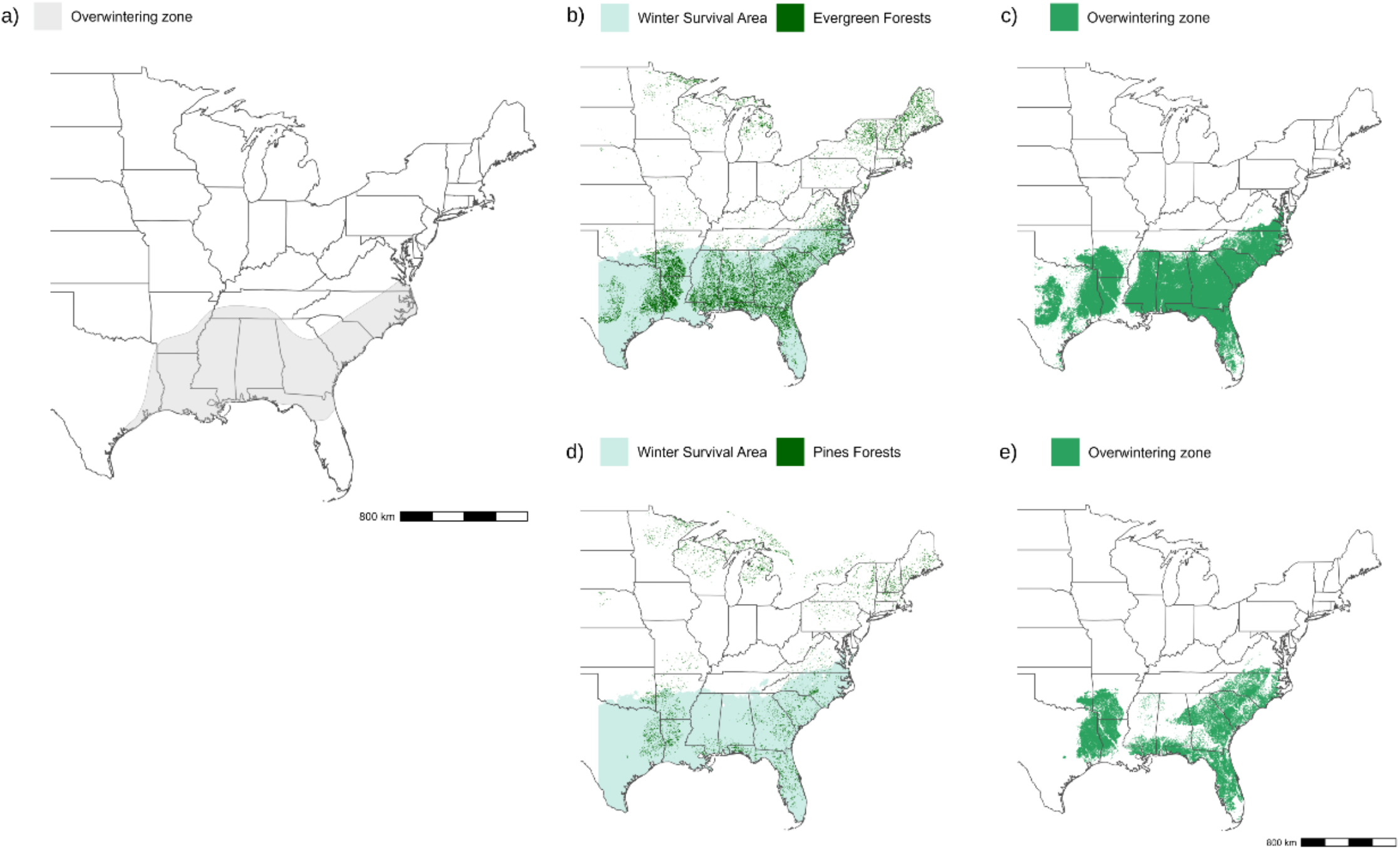
Estimated overwintering area for *Empoasca fabae*: (a) the estimate from Taylor and Shields (1995) (adapted), and the updated version based on data from 2024 for evergreen (b-c) and pines (d-e) forests. The winter survival zone is outlined by the common area estimated from 1900 to 2025 (white isoline in Figure 1b). The overwintering plots (c-e) were converted into polygons to improve visualization.

## DISCUSSION

The winter survival area of *E. fabae* has not expanded continuously over the past 126 years. Instead, it shows a non-trending pattern, with the Southeastern US remaining suitable over this period, while the northern range fluctuates latitudinally without a sustained shift. Even when analyzing the 1951–2012 interval (Baker et al., 2015), these results still showed no significant trends. Hence, the early spring arrival of *E. fabae* and its increased crop damage might be more parsimoniously explained by within-range warming, rather than by a continuous northward progression of the winter survival limits (Baker et al., 2015).

Warmer spring and mild winter temperatures accelerate the development of crop hosts (e.g., alfalfa, potato, and soybean) (Baker et al., 2015) and of woody perennials (Petrie et al., 2015) that *E. fabae* feeds (Lamp et al., 1994), respectively. Earlier hosts provide synchronized food resources, allowing populations to build sooner and reach damaging densities earlier, without requiring an expanded overwintering area (Baker et al., 2015). Because *E. fabae* development is temperature-dependent, warmer springs in the overwintering area shorten generation time, enabling earlier and potentially larger spring populations (Baker et al., 2015). Additionally, milder autumn conditions may increase *E. fabae* abundance later in the season (Lagos-Kutz et al., 2024) and delay the onset of diapause, though this last point remains underexplored.

Our updated estimate of the overwintering area broadly corresponds to that of Taylor & Shields (1995), yet it also reveals spatial variation influenced by the current distribution of winter hosts. Notably, Central Florida and the Texas Gulf Coastal Plain consistently appear suitable, whereas certain regions of Tennessee lack sufficient winter hosts. Importantly, the availability of these hosts is not static. Forest composition across the Southeastern US has undergone significant changes over the past century due to land use history, fire suppression, and reforestation (Hanberry, 2013). These shifts suggest that although the winter survival area remains relatively constant or even shifts farther north, the overwintering area may have contracted or expanded locally as evergreen and pine cover changed over time. Consequently, the overwintering area is dynamic both spatially and temporally, further complicating predictions of *E. fabae* overwintering population dynamics and underscoring the need for updated field data on winter host utilization (Taylor & Shields, 1995).

Our estimations used daily-average monthly minimum temperature rasters, but these may not capture short-term cold stress lasting only a few days that could lead to *E. fabae* complete mortality, especially when continuously exposed to −13°C (Decker & Maddox, 1967). We also assumed a fixed threshold of −9°C, as in Sidumo et al., (2005), but this does not account for microhabitat buffering provided by the hosts.

Our results indicate a dynamic winter survival area for *E. fabae*, suggesting that in some years this pest could overwinter farther north than in others, potentially leading to earlier local outbreaks without a permanent range shift. However, because these northern expansions are not sustained, they do not constitute a predictable, continuous expansion of the overwintering area. From a management perspective, this underscores the importance of annual monitoring to identify the source locations of migratory individuals arriving farther north within the potential overwintering area.

In conclusion, the winter survival area for *E. fabae* has not expanded continuously over the past 126 years. Rather, its current estimated overwintering area in the Southeastern US remains suitable, whereas the northern range varies dynamically. Future work should examine whether changes in spring synoptic wind patterns or extreme winter events influence interannual variability in the population’s northern extent.

## AUTHOR CONTRIBUTIONS

**Abraão Almeida Santos**: Conceptualization; data curation; formal analysis; investigation; methodology; visualization; writing – original draft; writing – review and editing. **Edel Pérez-López**: Resources; writing – review & editing.

## ACKNOWLEDGMENTS

We acknowledge the use of Grammarly (v. 1.2.250.1876) to assist with the language fluency and readability of this manuscript. All content was reviewed and approved by the authors, who take full responsibility for the accuracy and integrity of the work.

## DATA AVAILABILITY STATEMENT

All data supporting the findings of this study are available in the supporting information.

## SUPPORTING INFORMATION

**Supporting information S1:** Estimated winter survival area of *Empoasca fabae*.

